# A genetic mosaic mouse model illuminates the pre-malignant progression of basal-like breast cancer

**DOI:** 10.1101/2023.04.25.538333

**Authors:** Jianhao Zeng, Shambhavi Singh, Ying Jiang, Eli Casarez, Kristen A. Atkins, Kevin A. Janes, Hui Zong

**Affiliations:** Department of Microbiology, Immunology, and Cancer Biology, University of Virginia Health System, Charlottesville, VA 22908, USA; Department of Biomedical Engineering, University of Virginia, Charlottesville, VA 22908, USA; Department of Pathology, University of Virginia Health System, Charlottesville, VA 22908, USA; University of Virginia Comprehensive Cancer Center, University of Virginia Health System, Charlottesville, VA 22903, USA

**Keywords:** Basal-like breast cancer, *BRCA1*, pre-malignancy, genetic mosaic mouse model, spatiotemporal

## Abstract

Basal-like breast cancer is an aggressive breast cancer subtype, often characterized by a deficiency in *BRCA1* function and concomitant loss of *p53*. While conventional mouse models enable the investigation of its malignant stages, one that reveals its initiation and pre-malignant progression is lacking. Here, we leveraged a mouse genetic system known as <u>M</u>osaic <u>A</u>nalysis with <u>D</u>ouble <u>M</u>arkers (MADM) to generate rare GFP-labeled *Brca1*, *p53*-deficient cells alongside RFP+ wildtype sibling cells in the mammary gland. The mosaicism resembles the sporadic initiation of human cancer and enables spatially resolved analysis of mutant cells in comparison to paired wildtype sibling cells. Mammary tumors arising in the model show transcriptomic and genomic characteristics similar to human basal-like breast cancer. Analysis of GFP+ mutant cells at interval time points before malignancy revealed a stepwise progression of lesions from focal expansion to hyper-alveolarization and then to micro-invasion. These stereotyped morphologies indicate the pre-malignant stage irrespective of the time point at which it is observed. Paired analysis of GFP-RFP siblings during focal expansion suggested that hyper-alveolarized structures originate from ductal rather than alveolar cells, despite their morphological similarities to alveoli. Evidence for luminal-to-basal transition at the pre-malignant stages was restricted to cells that had escaped hyper-alveoli and progressed to micro-invasive lesions. Our MADM-based mouse model presents a useful tool for studying the pre-malignancy of basal-like breast cancer.

**Summary statement:** A mouse model recapitulates the process of human basal-like breast tumorigenesis initiated from sporadic *Brca1, p53*-deficient cells, empowering spatially-resolved analysis of mutant cells during pre-malignant progression.

## Introduction

Breast cancer is the most frequently diagnosed cancer type and the second leading cause of cancer death in women (Siegel et al., 2022). Human breast cancer is a heterogeneous disease classified into six molecular subtypes with distinct prognosis: luminal A, luminal B, HER2-enriched, normal-like, claudin-low, and basal-like (Perou et al., 2000; Prat et al., 2010; Tobin et al., 2015). Basal-like breast cancer accounts for 15–20% of breast cancer cases and is the most aggressive subtype with earlier onset, increased chance of metastasis, and absence of hormonal-therapy targets (Fulford et al., 2007; Millikan et al., 2008; Tobin et al., 2015; Turner et al., 2004). Basal-like breast cancers show a high prevalence of *p53* mutations (∼80%) and deficiency in homology-directed DNA repair (∼50%); the latter is often caused by germline mutations in *BRCA1/2*, somatic epigenetic inactivation of *BRCA1/2*, or the loss of other essential genes for homology-directed DNA repair pathway (known as “BRCAness”) (Lord and Ashworth, 2016; McCabe et al., 2006; Cancer Genome Atlas Network, 2012; Tian et al., 2019). Early detection and prevention of basal-like breast cancer can fundamentally improve patient care, particularly for individuals with germline *BRCA1* mutations who are at a high risk of breast cancer. Therefore, it is imperative to gain a deep understanding of how basal-like breast cancer initiates and progresses during the pre-malignant stages.

Genetically engineered mouse models (GEMMs) present invaluable pre-clinical resources for studying human basal-like breast cancer. Conditional knockout of *Brca1* and *p53* in mouse mammary epithelial cells gives rise to mammary tumors resembling human basal-like breast cancer (Hollern et al., 2019; Liu et al., 2007; Molyneux et al., 2010; Xu et al., 1999). These models are useful for studying cancer at the malignant stage; however, examining cancer initiation and pre-malignant progression with these GEMMs faces considerable limitations. First, conditional knockout models generate numerous rather than rare mutant cells at the cancer initiation stage; thus, they do not accurately mimic human cancer initiation from sporadic mutant cells (Liu et al., 2011; Muzumdar et al., 2007), which may impact pre-malignant development. Second, even if rare mutant cells can be generated, unequivocally pinpointing subtle aberrant behaviors of *BRCA1* mutant cells at the pre-malignant stage remains challenging.

To overcome these limitations, we used a mouse genetic system known as Mosaic Analysis with Double Markers (MADM) developed by our lab (**Fig. 1A**). MADM consists of a pair of chimeric GFP and RFP coding sequences (separated by a loxP-containing intron) knocked into homologous chromosomes. Each knock-in cassette is syntenic with either the wildtype or mutant allele of one or more tumor suppressor genes on the same chromosome. From a non-labeled heterozygous animal, Cre/loxP-mediated inter-chromosomal mitotic recombination followed by X-segregation of chromosomes generates a homozygous mutant cell labeled with GFP and its sibling wildtype cell labeled with RFP. Mutant cells are rare (0.1%–1% or even lower) due to the low frequency of inter-chromosomal recombination (Muzumdar et al., 2007; Zong et al., 2005), thereby approximating sporadic cancer initiation. The permanent GFP labeling of rare mutant cells enables spatially resolved tracking of their behavior at any time point during tumorigenesis (**Fig. 1A)** (Liu et al., 2011; Yao et al., 2020; Zong et al., 2005). Furthermore, along with a GFP+ mutant cell, MADM generates a sibling RFP+ wildtype cell simultaneously, which serves as a perfect internal reference enabling the detection of subtle abnormalities of GFP+ mutant cells. Here, we applied MADM to engineer a mosaic genetic mouse model for basal-like breast cancer, in which cancer initiates with sparse *Brca1, p53*-deficient cells in the mammary gland. Delineating GFP+ mutant cells up to early malignancy yields insights into cancer initiation and pre-malignant progression, creating future opportunities for deeper analysis and tests of preventative interventions.

**Fig. 1.**
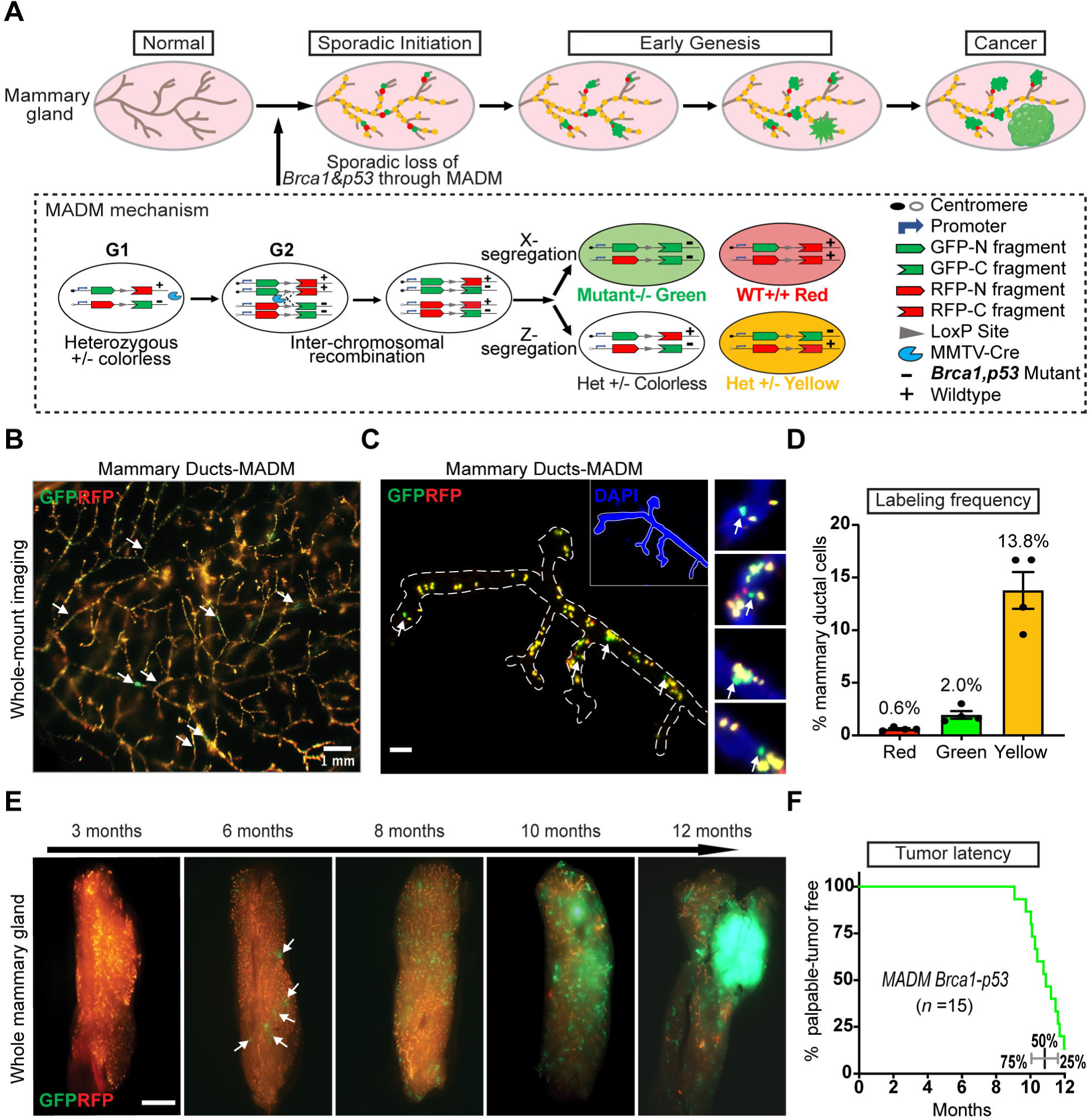
A MADM model that tracks the process of *Brca1, p53*-dependent mammary tumorigenesis from sporadic cancer-initiating cells to frank tumors. (A) Modeling breast cancer development from sporadic mutant cells to tumors with MADM. From a colorless heterozygous animal, through inter-chromosomal recombination in mitotic cells at the G2 phase, MADM generates one GFP+ mutant cell and one RFP+ wildtype cells after X segregation (two recombinant sister chromatids segregate into different daughter cells); alternatively, Z segregation generates one colorless and one dual-colored (yellow) cell that are both heterozygous (two recombinant sister chromatids segregate into the same daughter cells). The schematic of MADM is reproduced with permission from ref. (Zong et al., 2005), Cell Press. (B) The MADM model induces sparse and scattered GFP+ mutant cells (arrows) in mouse mammary glands. The image is representative of four MADM-mutant mice collected at three months old for whole-mount fluorescence imaging. Scale bar =1 mm. (C) High-resolution imaging of the GFP+ mutant cells (arrows) in mammary glands from three-month- old MADM-mutant mice. The image is representative of mammary tissue sections acquired from four mice at three months old and imaged by wide-field fluorescence microscopy. Scale bar =100 μm. (D) The proportion of MADM-labeled cells among all mammary epithelial cells in MADM-mutant mice at three months old. Sections of mammary glands were imaged with wide-field microscopy. The number of MADM-colored and mammary ductal cells was counted by GFP/RFP fluorescence and DAPI, respectively. Data are shown as the mean percentage ± s.d. from *n* =4 mice. (E) Fluorescence imaging of whole mammary glands from a cohort of MADM-mutant mice at different ages, showing the progressive expansion of GFP+ foci and tumor formation. Arrow indicates GFP+ focal expansions. Images were collected on a fluorescence stereomicroscope and are representative of 10 mice for each age. Scale bar =500 μm. (F) Percent tumor-free of MADM-mutant mice (*n* =15) after 12 months as the endpoint, showing a median latency of 11 months. Quantiles at the bottom show a narrow spreading of tumor latency.

## Results

### MADM model reveals the process of mammary tumorigenesis initiated by sporadic loss of *Brca1* and *p53*

To establish a MADM-based mouse model of breast cancer, we prepared two stock mouse lines through a multi-generational breeding scheme (**Fig. S1)**. For one stock line, we bred the *Brca1* and *p53* mutant alleles (Jacks et al., 1994; Xu et al., 1999) as cancer-initiating mutations onto the MADM-TG line (Hippenmeyer et al., 2010). For the other stock line, we introduced the *MMTV-Cre* transgene (Wagner et al., 2001) into the MADM-GT line (Hippenmeyer et al., 2010) to target mammary epithelial cells broadly. Inter-crossing between the two stock lines generates MADM*-Brca1*-*p53; MMTV-Cre* (MADM-mutant) mice (**Fig. S1B**), in which Cre-mediated inter-chromosomal recombination during G2 phase and mitosis generates sparse GFP+ homozygous *Brca1* and *p53* mutant cells predisposed to become cancerous (**Fig. 1A)**.

The rarity of initiated GFP+ mutants in MADM is more reflective of human cancer initiation and enables clonal analysis of pre-malignant expansion (Greaves and Maley, 2012; Knudson, 1971; Muzumdar et al., 2007). We assessed the abundance of GFP+ mutant cells in mammary glands from MADM-mutant mice at three months old, an age shortly after the peak of *MMTV-Cre* expression (Buono et al., 2006; Wagner et al., 1997). Using whole-mount fluorescence imaging of mammary glands, we found a high abundance of heterozygous yellow cells (GFP and RFP double positive) generated from MADM recombination events in G1 or post-mitotic cells (G0) (**Fig. S2A)**. These non-mutant controls illuminate the mammary ductal system and confirm in vivo recombination by *MMTV-Cre*. Among many yellow cells, we observed sparse and scattered GFP+ mutant cells along mammary ducts (**Fig. 1B)**. For a higher resolution view, we sectioned mammary glands and performed confocal imaging to visualize the abundance of GFP+ mutant cells. We found the GFP+ mutant cells were often singular (**Fig. 1C)** and accounted for ∼2% of all mammary epithelial cells (**Fig. 1D)**. The proportion of RFP+ wildtype sibling cells was lower (∼0.6%) than that of GFP+ mutant cells, indicating a possible early survival disadvantage of the wildtype cells compared to the mutants. Notably, almost all MADM-labeled cells were positive for cytokeratin 8 (CK8, marker for mammary luminal cells) but negative for cytokeratin 14 (CK14, marker for mammary basal cells) (**Fig. S2B, C),** suggesting that they arose from the luminal layer where the reported cell of origin for basal-like breast cancer resides (Lim et al., 2009; Molyneux et al., 2010; Shehata et al., 2012).

To assess the progression of initiated mutants, we collected mammary glands from MADM-mutant mice at different time points (10 mice at each age) and evaluated the overall expansion of GFP+ cells through whole-mount imaging. At 3 months, the expansion of GFP+ mutant cells was barely noticeable, but from 6 months onward, multicellular GFP+ foci broadly expanded and gradually progressed to GFP+ tumors (**Fig. 1E)**. By examining a cohort of 15 mice with an endpoint of 12 months of age, we found 13 (87%) developed GFP+ tumors with a median latency of 11 months (**Fig. 1F)**. Despite the rarity of cancer-initiating mutant cells, the MADM-mutant mice show a tumor latency only slightly longer than the ∼ 9 months latency of conditional knockout mouse model that initiates cancer with numerous mutant cells (Xu et al., 1999). This suggests that the abundance of cancer-initiating cells is not a rate-limiting factor for the kinetics of breast cancers driven by *Brca1* and *p53* deficiency.

### MADM-mutant mammary tumors resemble human basal-like breast cancer

Human basal-like breast cancers are characterized by a high proliferation index, lack of estrogen receptor (ER), progesterone receptor (PR), and HER2 over-expression (Palacios et al., 2005; Perou et al., 2000; Rakha et al., 2008). We assessed 6 MADM tumors of their proliferation and hormone receptor expression by immunohistochemistry and found them highly positive for Ki67 (∼70% cells) and mostly negative for ER, PR, and HER2 **(Fig. 2A, S3A)**, matching the histopathological features of human basal-like tumors. To further examine whether the MADM tumors resemble human basal-like breast cancer at the molecular level, we performed RNA sequencing of 12 MADM tumors and extracted a panel of 50 genes (PAM50) used to stratify breast tumor subtypes (Perou, 2011). We co-clustered PAM50 signatures from our MADM tumors with the profiles of 1104 human breast tumors from TCGA annotated for five breast cancer subtypes. To mitigate overall differences in gene abundance across species, we identified a unique set of mouse-to-human orthologs across all TCGA and MADM tumors and normalized each sample to obtain relative expression values for each gene (see methods for details). We found MADM tumors clustered with the human basal-like subtype but not others **(Fig. 2B)**, suggesting close similarity. Another hallmark of human *BRCA1-*mutated basal-like breast cancer is the high frequency of copy number variations (CNVs) for genomic loci containing oncogenes or tumor suppressors (Annunziato et al., 2019; Weigman et al., 2012). To determine whether MADM tumors also share this hallmark, we conducted whole-exome sequencing on six MADM tumors and paired normal somatic tissue. We found recurrent amplification of multiple chromosomal segments harboring oncogenes—such as *Met*, *Myc,* and *Fgfr1*—along with recurrent deletion of the tumor suppressor gene *Rb1* **(Fig. 2C)**. These findings correspond closely with frequent copy number variations or transcriptional changes observed in human basal-like tumors **(Fig. S3B)** (Cancer Genome Atlas Network, 2012). Collectively, the histopathological, transcriptomic, and genomic analyses of MADM tumors demonstrate that our MADM-mutant mice represent an authentic model for human basal-like breast cancer.

**Fig. 2.**
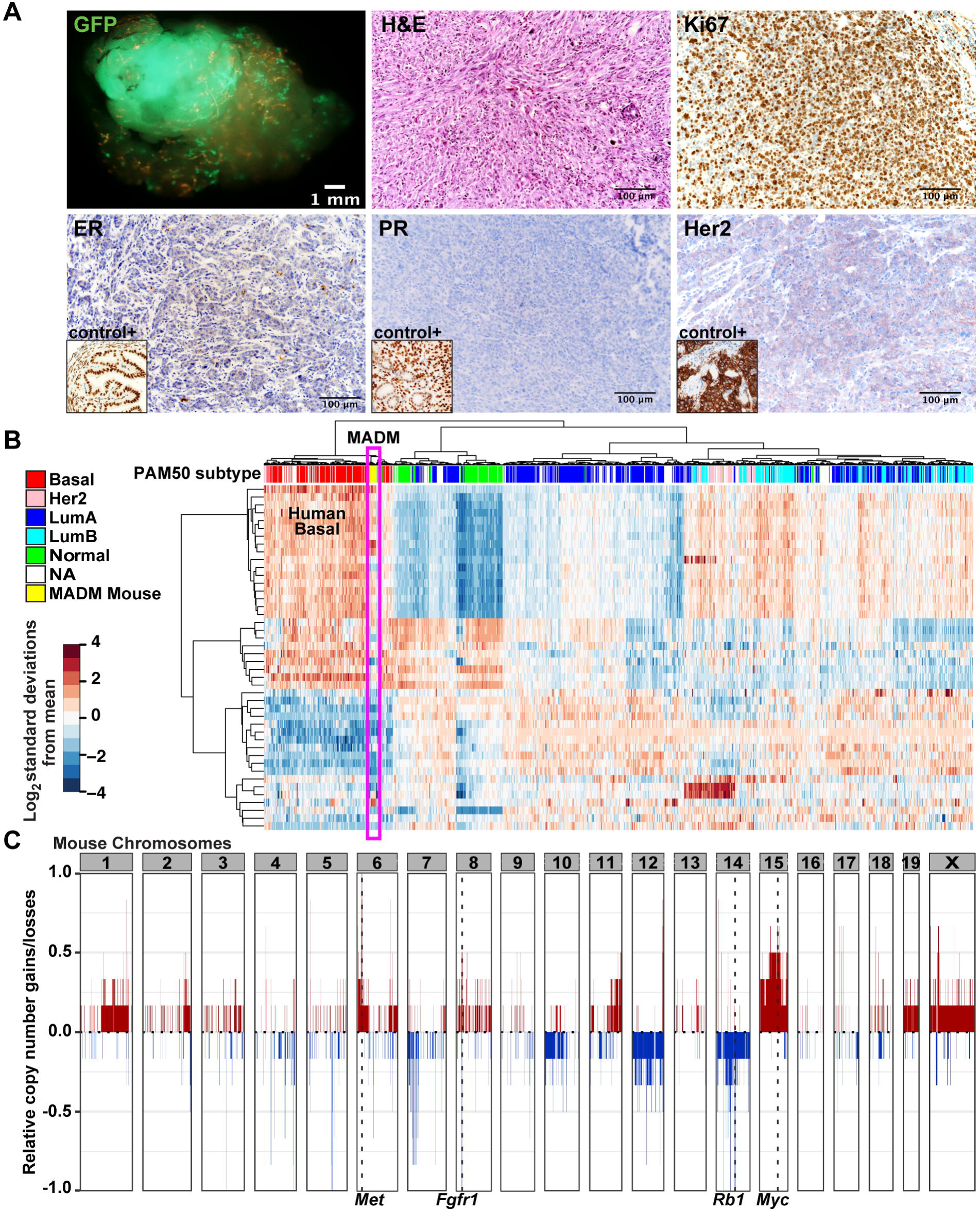
MADM-mutant mammary tumors resemble human basal-like breast cancer. (A) Whole-mount fluorescence image of GFP+ mammary tumors (upper left); H&E (upper middle), immunohistochemistry staining of Ki67 (upper right), estrogen receptor (ER, lower left with a mouse oviduct as a positive control in the inset), progesterone receptor (PR, lower middle with a mouse uterus as a positive control in the inset), and Her2 (lower right with a mouse HER2-amplified mammary tumor as a positive control in the inset). Representative images of tumors from 6 mice. Scale bar =100 μm. (B) PAM50-based clustering of MADM tumors with human breast cancers previously subtyped by PAM50 analysis. 12 MADM tumors and 1104 human breast tumors from TCGA datasets were analyzed. Cross-species differences were normalized using a set of mouse-to-human orthologs (see methods for details). (C) Copy-number variations (CNVs) in six MADM tumors were assessed by whole-exome sequencing. Gains in *Met, Fgfr1,* and *Myc*, and loss of *Rb1* are highlighted.

### MADM resolves the characteristic morphological stages of pre-malignant progression

After confirming sparse initiation and progression to full-blown tumors, we next investigated the successive phenotypic alterations of MADM mutant cells throughout pre-malignancy. We harvested mammary glands from a cohort of mice at the intermediate ages between cancer initiation and tumor formation; then, we performed tissue clearing with CUBIC (Susaki et al., 2015) and conducted whole-mount, high-resolution 3D imaging using light-sheet microscopy **(Fig. S4A)**. We detected stepwise morphological abnormalities by comparing green mutant ducts with yellow heterozygous ducts **(Fig. 3A)**. At 3 months old, we observed stretches of mutant cells that occupied a continuous region of the mammary duct without noticeable alterations in ductal morphology. At 6 months old, some GFP+ mutant cells extended side branches resulting in a slightly more complex morphology than yellow ducts. This hyper-proliferation of mutant cells preceding prominent changes in tissue organization is consistent with observations in *BRCA1*-mutant carriers (Martins et al., 2012; McKian et al., 2009). Upon further expansion at 8 months, some mutant branches developed extensive epithelial buds reminiscent of alveologenesis during early pregnancy (Richert et al., 2000), which was notable for virgin MADM-mutant females. The alveologenesis was pervasive in late-stage mammary glands and exhibited distinct histology when compared with the internal control yellow heterozygous ducts (**Fig. S4B, C)**. We further quantified the number of buds (alveoli) per 100-µm primary ducts and found that mutant ducts had over ten-fold more alveoli than the controls **(Fig. S4D, E)**. Binning the data into four levels based on the number of alveoli per 100 µm primary ducts—level 0 (<5), level I (5-20), level II (20-50), and level III (>50) **(Fig. S4F)**—we found that high-level alveologenesis (II-III) exclusively occurred within the mutant ducts **(Fig. S4G)**, suggesting that this “hyper-alveolarization” is a characteristic phase of early malignancy. In later 10-month-old mice approaching tumor onset, we occasionally observed tiny GFP+ spherical masses (<1 mm in diameter) that had lost all ductal morphology but were not yet palpable (**Fig. 3A)**. These spherical outgrowths of GFP+ mutant cells into the stroma represent the onset of malignancy and were classified as “micro-invasion” hereafter. Often, mutant ducts showed various expansion levels at each age, with the morphological hallmarks of hyper-alveolarization and micro-invasion representing the most extensive clonal expansion at each age **(Fig. S4B)**.

**Fig. 3.**
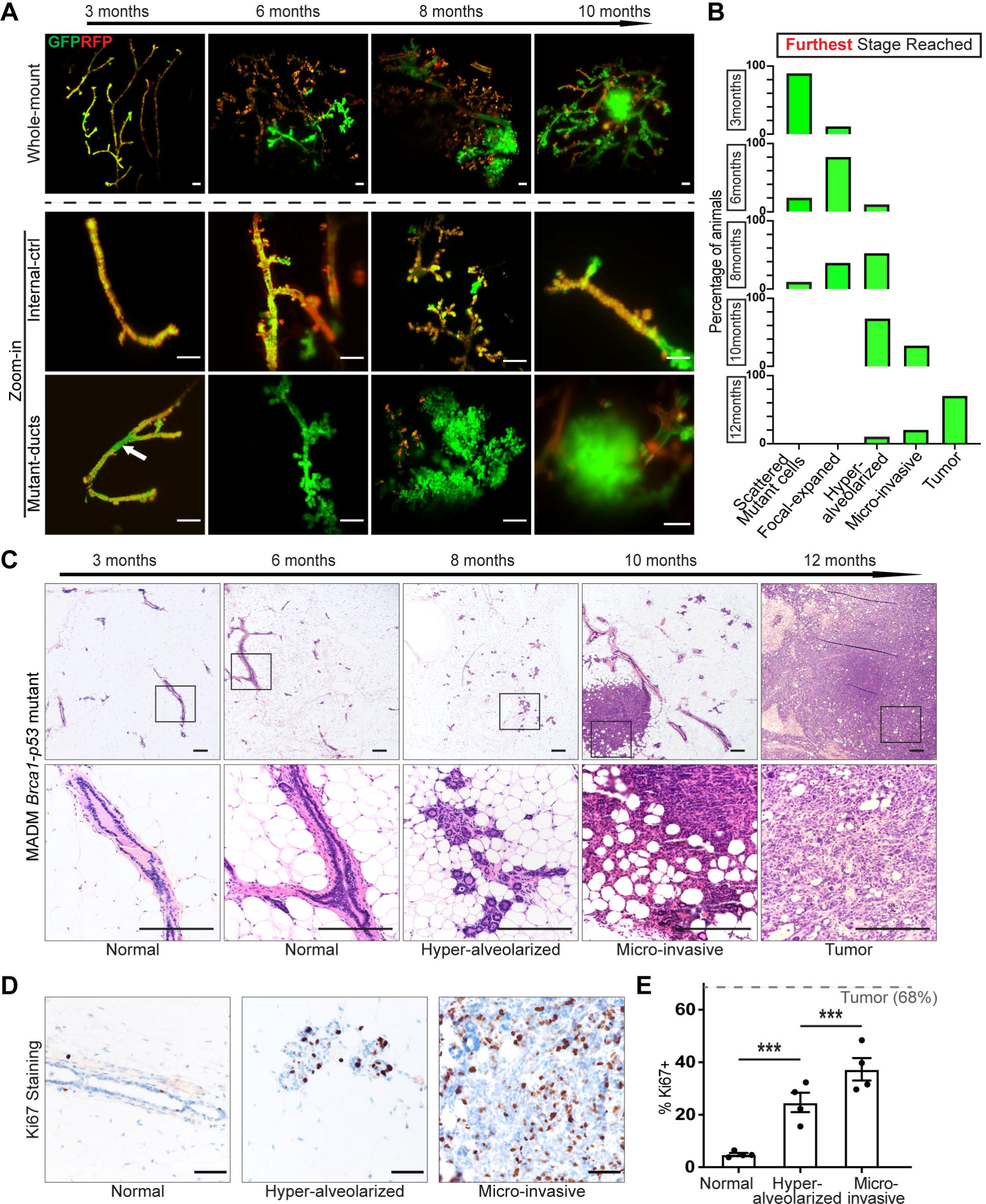
MADM-mutant mammary glands resolve stereotyped changes of basal-like premalignancy. (A) Progressive morphological changes in green mutant ducts compared to yellow internal control heterozygous ducts as mice age. Upper panel: Low-magnification 3D fluorescence imaging of mammary glands from a cohort of mice at different ages; Middle and lower panel: Higher magnification of the control yellow ducts and the green mutant ducts. Arrow in the left panel indicates a GFP+ mutant region within the mammary ducts. Mammary glands from MADM-mutant mice at each age (*n* =10) were cleared with the CUBIC method, and 3D high-resolution images were acquired by light-sheet microscopy. Scale bar =100 μm. (B) Timeline of the first occurrence of each pre-malignant stage of GFP+ mutant foci. The morphology of all mutant ducts in MADM-mutant mice at each age (*n* =10) was examined by whole-mammary-gland fluorescence imaging. Mice are categorized based on the furthest stage of premalignancy reached. The proportional distribution of mice at each age is plotted. (C) H&E staining of the progressive change of mammary ductal shape from normal to hyper-alveolarized, to micro-invasive, and to tumors. Paraffin slides of mammary glands from MADM-mutant mice at each age (*n* =10) were used for staining. Scale bar =200 μm. (D) Ki67 staining of normal mammary ducts and MADM-mutant lesions at different stages. Ducts of each morphology were collected from four mice. Scale bar =100 μm. (E) Quantification of Ki67 percent positivity (Ki67+%) of cells within normal ducts and MADM-mutant lesions at the hyper-alveolarized and micro-invasive stages. The dashed line indicates Ki67+% in frank tumors (Fig. 2A). Data are represented as mean ± s.e.m. from *n* =4 mice. ***<0.001 by Mann–Whitney test.

To further determine whether the hyper-alveolarized ducts and the micro-invasions reflect a sequential progression of mammary cancer in MADM-mutant animals, we evaluated whether there is a temporal sequence in the occurrence of these structures. Classifying the most-advanced GFP+ lesion observed in the mammary gland of each MADM-mutant animal at a given time point, we observed a sequential emergence of focally expanded GFP+ cells, hyper-alveolarized GFP+ ducts, and micro-invasions, culminating in the emergence of GFP+ tumors at one year (**Fig. 3B).** Histological analysis further supported a temporal sequence of these pre-malignant structures. While mammary epithelial cells from control wildtype mice appeared normal across all ages **(Fig. S5A)**, mutant cells in hyper-alveolarized ducts prominent at 8 months displayed abnormal nucleomegaly (about 1.5x larger than wildtype nuclei), small but conspicuous nucleoli, and loss of the basally-oriented nuclear polarity **(Fig. 3C, S5B)**. The micro-invasions contained residual hyper-alveolarized lobular units along with more advanced micro-invasive carcinoma (measuring less than 1 mm) comprised of individual cells and larger cords that elicited a stromal response **(Fig. 3C)**. Finally, the frank tumors were dominated by infiltrating cells with a desmoplastic stromal response; no remnants of alveoli were visible. The progressive transition from hyper-alveolarization to micro-invasion was further supported by Ki67 staining of each phase, revealing a gradual increase in cell proliferation toward levels observed in fully developed tumors **(Fig. 3D, E)**. Despite a virtually nonexistent carcinoma in situ phase, the MADM-mutant model suggests that cancer-initiating cells progress through a visually identifiable sequence of ductal morphologies before the emergence of mammary tumors.

### Hyper-alveolarized structures arise from mutant ductal cells

The mammary epithelium is composed of two anatomically and functionally distinct compartments—the alveolar regions that produce milk during lactation and the ductal regions that drain milk to the nipple (**Fig. 4A)** (Visvader, 2009; Visvader and Stingl, 2014). It was unclear whether the hyper-alveolarized structures consisting of mutant cells originated from alveolar or ductal mutants. Since MADM generates one RFP+ wildtype sibling cell alongside each original GFP+ mutant cell (**Fig. 1A)**, we investigated this question with a “twin-spot” analysis that evaluates the expansion of mutant clones compared to wildtype sibling clones within ductal and alveolar compartments, respectively (Espinosa and Luo, 2008; Muzumdar et al., 2007; Terry et al., 2020).

**Fig. 4.**
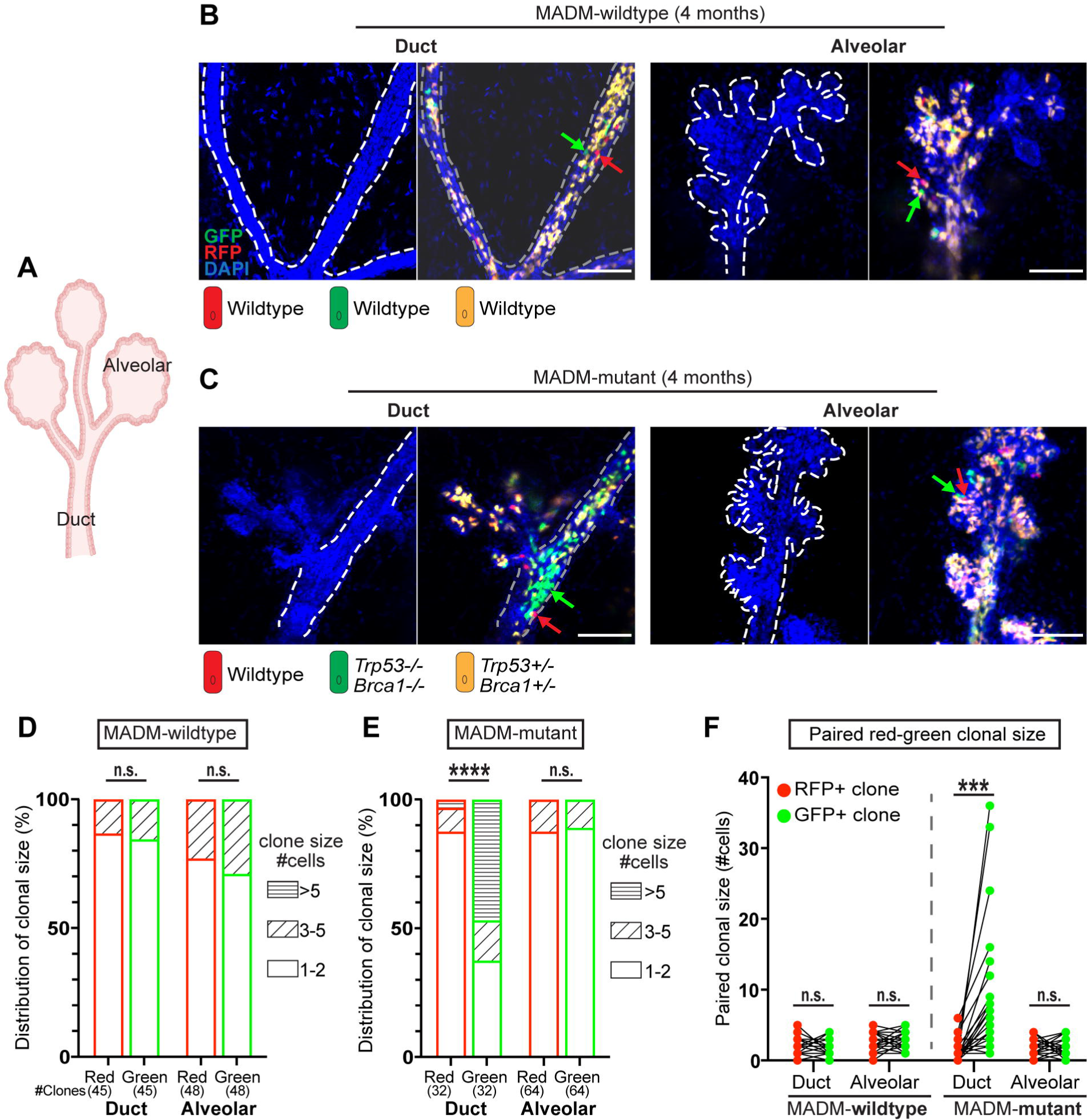
Ductal but not alveolar cells show initial clonal expansion in MADM-mutant glands. (A) Schematic of ductal and alveolar regions in the mouse mammary gland. (B) GFP+ and RFP+ clone pairs of MADM-wildtype mice are of similar size and do not show prominent expansion. 3D fluorescence images of cleared mammary glands from MADM-wildtype mice at four months old (*n*=4) were acquired as Z-stacks on a confocal microscope. The green arrows indicate the GFP+ cells, and the red arrows indicate RFP+ cells. Scale bar =100 μm. (C) GFP+ mutant clones in the ductal region of MADM-mutant mice show prominent expansion, while their sibling RFP+ clones do not. In the alveolar region, neither GFP+ nor RFP+ clones expand. 3D fluorescence images of cleared mammary glands from MADM-mutant mice at four months old (*n*=4) were acquired as Z-stacks on a confocal microscope. The green arrows indicate the GFP+ cells, and the red arrows indicate RFP+ cells. Scale bar =100 μm. (D) Size distribution of GFP+ and RFP+ clones in ductal and alveolar regions of mammary glands from MADM-wildtype mice at four months old. Data were pooled from four mice, and the total number of clones is indicated for each group. n.s. p>0.05 by Fisher’s Exact test. (E) The clonal size distribution of GFP+ and RFP+ clones in ductal and alveolar regions of mammary glands from MADM-mutant mice at four months old. Data were pooled from four mice, and the total number of clones is indicated for each group. n.s. p>0.05, ***p<0.0001 by Fisher’s Exact test. (F) Quantification of the clonal size for paired GFP+ and RFP+ sibling clones in the ductal and alveolar regions of mammary glands from both MADM-wildtype and MADM-mutant mice at four months. Clones from four mice were pooled. Data are represented as mean ± s.e.m. ***p<0.001 by paired t-test.

For the twin-spot analysis, we selected mice at four months old because the focal expansion of GFP+ mutant cells was evident at this age, while hyper-alveologenesis was not yet present (**Fig. 3B)**. As a reference, we initially analyzed 40 twin spots in the ductal or alveolar regions of four MADM-wildtype mice that lacked *Brca1* and *p53* mutant alleles (**Fig. S1B)**; thus, both green and red cells are wildtype. In MADM-wildtype animals, GFP+/RFP+ clones did not expand prominently in ducts or alveoli **(Fig. 4B)**, and quantitative analysis revealed no overall bias in expansion for either compartment **(Fig. 4D)**. In contrast, when we performed the same analysis with four MADM-mutant mice (GFP+: mutant; RFP+: wildtype), the ductal region showed prominent expansion of GFP+ mutant clones over RFP+ wildtype clones (**Fig. 4C, E)**. Surprisingly, the mutant clones in alveoli, even though harboring the same *Brca1* and *p53* mutations as those in ducts, neither expanded nor differed in size from the neighboring RFP+ wildtype clones. To conduct the twin-spot analysis more rigorously, we compared the sizes of the GFP+ and RFP+ sibling clones in a strictly pairwise manner. Our analysis revealed that only GFP+ mutant clones within the ductal regions exhibited significantly larger sizes than their RFP+ sibling clones (**Fig. 4F)**, indicating that the initial clonal expansion of *Brca1, p53*-mutated mammary cancer occurs exclusively in the ductal region. While multiple studies on human tissues and mouse models have described the outgrowth of alveolar-like buds during the pre-malignant development of breast cancer and suggested aberrant alveolar cell expansion (Bach et al., 2021; McKian et al., 2009; Poole et al., 2006; Tao et al., 2017), our results raise the possibility that such outgrowth has a ductal rather than an alveolar origin, despite their morphological resemblance to the latter.

### The onset of luminal-to-basal transition coincides with the appearance of micro-invasion

Basal-like breast cancers express basal-cell markers (CK5/14, α-SMA, P63, etc.) yet arise from luminal progenitor cells (Lim et al., 2009; Liu et al., 2007; Molyneux et al., 2010; Shehata et al., 2012). This luminal-to-basal transition is thought to be critical for cancer progression, as it promotes stemness and invasiveness of *BRCA1*- or *p53*-mutant mammary epithelial cells in vitro (Bai et al., 2022; Kim et al., 2011; Liu et al., 2008; Mizuno et al., 2010). However, in vivo mapping of luminal-to-basal transitions during tumorigenesis is lacking. To determine whether luminal-to-basal transition occurs in the two representative pre-malignant stages of MADM-mutant mice (hyper-alveolarized ducts and micro-invasions), we leveraged the single-cell resolution of MADM and assessed the morphology of mutant cells at each stage. Normal mammary luminal cells present a cuboidal shape, whereas basal cells show an elongated spindle shape (Rios et al., 2014). Within hyper-alveolarized ducts, mutant cells mostly exhibited a cuboidal shape with relatively homogeneous cell size **(Fig. 5A upper panel)**. In contrast, mutant cells that were micro-invasive showed heterogeneous morphologies, with a fraction adopting an elongated spindle shape reminiscent of basal cells **(Fig. 5A lower panel)**. We quantified the cell size and circularity of mutant cells in hyper-alveolarized ducts and micro-invasions from four MADM-mutant mice each. Overall, the mutant cells in micro-invasions were significantly larger in size and lower in circularity than those in hyper-alveolarized ducts **(Fig. 5B, C)**, implying a transition of cell state between these two stages.

**Fig. 5.**
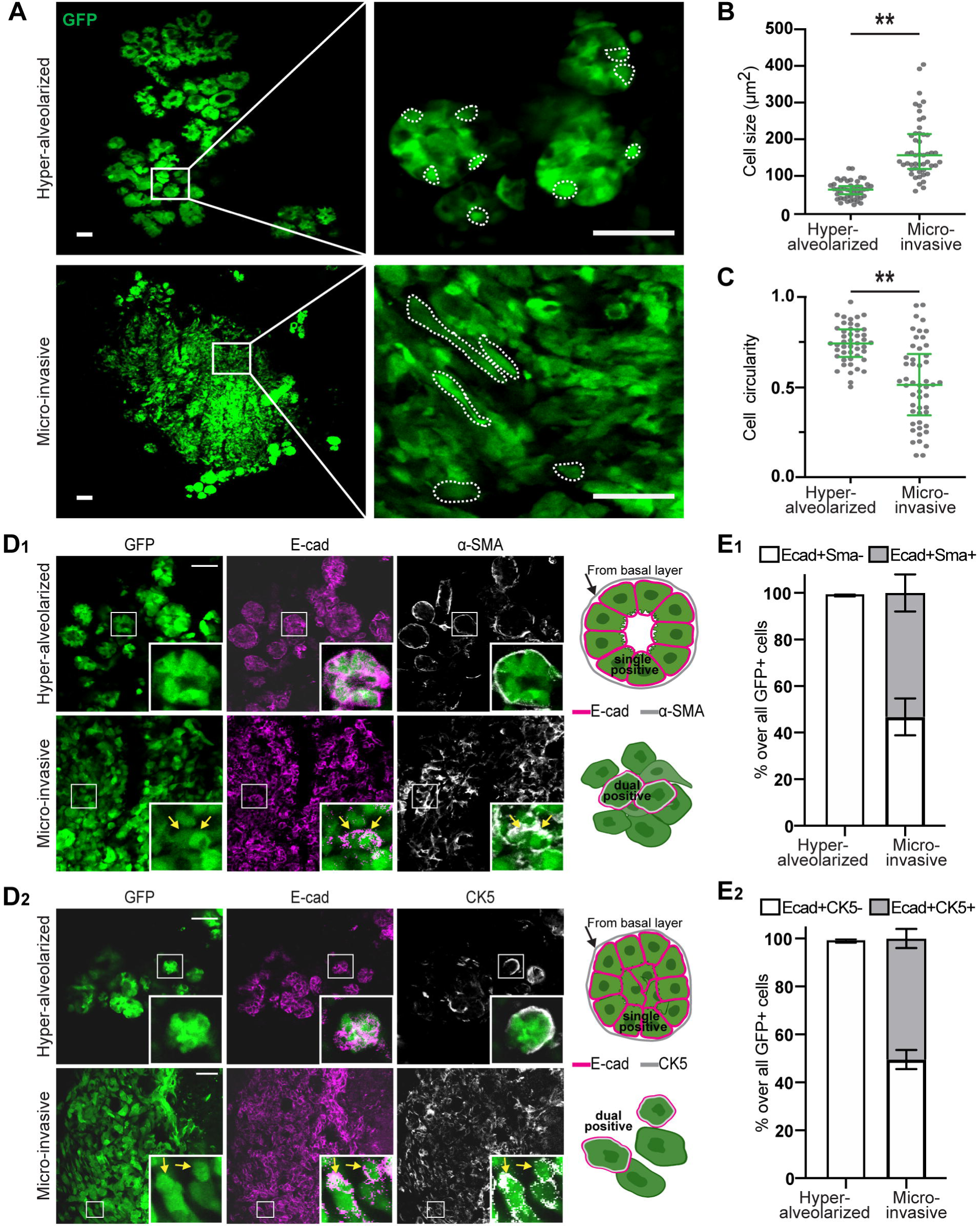
MADM-mutant cells undergo a partial luminal-to-basal transition upon micro-invasion. (A) Morphology of GFP+ mutant cells in hyper-alveolarized ducts and micro-invasive lesions. Mammary gland sections were imaged by confocal microscopy. For each feature, samples from four mice were assessed. The dashed regions delineate single cells. Scale bar =50 μm. (B) Size of mutant cells in hyper-alveolarized ducts and micro-invasions. Cells from four mice were pooled. Data are represented as median ± interquartile range from *n* =50 cells. **<0.01, by nested t-test. (C) The circularity of mutant cells in hyper-alveolarized ducts and micro-invasions. Cells from four mice were pooled. Quartiles are shown. Data are represented as median ± interquartile range from *n* =50 cells. **<0.01, by nested t-test. (D) In hyper-alveolarized ducts, GFP+ mutant cells were exclusively positive for E-cadherin but negative (outside of the GFP+ signal) for α-SMA(**D_1_**) or CK5 (**D_2_**), whereas in micro-invasions, some mutant cells were E-cad+ α-SMA+ dual positive(**D_1_**) or E-cad+CK5+ dual positive (**D_2_**). Frozen sections of mammary glands were stained and imaged by confocal microscopy. For each feature, samples from four mice were analyzed. Scale bar =50 μm. (E) E_1_: the proportion of E-cad+SMA-(luminal) and E-cad+SMA+ (partially transitioned) mutant cells; E_2_ the proportion of E-cad+CK5-(luminal) and E-cad+CK5+ (partially transitioned) mutant cells; in hyper-alveolarized ducts and “micro-invasions. For each feature, a total of ∼700 mutant cells from mammary glands from four mice were assessed. Data are represented as mean ± s.d.

To clarify whether the morphological change represents a luminal-to-basal transition, we assessed the expression of a luminal cell marker (E-Cadherin) and two basal cell markers (α-SMA, CK5) in mutant cells by immunofluorescent staining. In hyper-alveolarized ducts, GFP+ mutant cells maintained the expression of E-cadherin without acquiring basal cell markers that are present in the myoepithelial cells surrounding GFP+ luminal cells **(Fig. 5D_1_, D_2_ upper panel).** In micro-invasions, by contrast, mutant cells often jointly expressed E-cadherin and multiple basal cell markers **(Fig. 5D_1_, D_2_ lower panel),** suggesting a partial transition from the luminal to basal cell state. We quantified this observation by analyzing a total of ∼700 cells in mammary tissues from four mice with hyper-alveolarized ducts or micro-invasions. In hyper-alveolarized ducts, E-Cadherin-positive mutant cells were almost exclusively negative for α-SMA and CK5, but ∼50% of mutant cells expressed both basal cell markers and E-Cadherin in micro-invasions **(Fig. 5E_1_, E_2_)**, suggesting a hybrid cell state of future interest for molecular analysis and possibly therapeutic targeting.

## Discussion

In this study, we established a genetic mosaic mouse model for basal-like breast cancer, validated by its histopathological, transcriptomic, and genomic similarities to human basal-like breast cancers. Taking advantage of the spatial resolution provided by MADM, we identified multiple morphologically distinct pre-malignant stages, including focal expansion of mutant cells, hyper-alveolarization of mutant ducts, and micro-invasion. Surprisingly, the clonal analysis revealed that hyper-alveolarized mutant structures originate from ductal rather than alveolar cells. Further progression from hyper-alveolarized structures to micro-invasions results in loss of ductal organization and a partial luminal-to-basal transition, manifested by enlarged cell size, elongated cell shape, and elevated cell proliferation. Taken together, Our MADM-based mouse model presents a useful tool for studying the pre-malignancy of basal-like breast cancer with an excellent spatial resolution, which should empower pre-clinical research on early detection and cancer prevention.

Cancer modeling by MADM offers several unique advantages. Firstly, MADM-mutant mice create a small number of homozygous mutant cells from heterozygous mother cells, which is reminiscent of sporadic loss of heterozygosity in cancer. For germline *BRCA1* mutation carriers, cancer is often initiated by the sporadic loss of the wildtype allele (Cornelis et al., 1995; Maxwell et al., 2017; Nones et al., 2019). Secondly, the generation of mutant cells is tightly coupled with permanent GFP labeling in a single mitotic recombination event, allowing mutant cells to be examined throughout the entire process of tumorigenesis (Muzumdar et al., 2007; Zong et al., 2005). MADM fluorescence labels are bright enough to be visualized in fresh, intact glands under a fluorescence stereomicroscope (**Fig. 1E**), allowing for the gross evaluation of pre-malignant status before various downstream analyses that require unfixed tissue or live cells. Thirdly, MADM generates sibling wildtype cells that are labeled with RFP, providing an internal control for the GFP-labeled mutant cells (Beattie et al., 2017; Terry et al., 2020). Without RFP+ wildtype siblings as a reference **(Fig. 4)**, it would be difficult to distinguish the clonal expansion of GFP+ mutant ductal cells from stochastic neutral drift (Lopez-Garcia et al., 2010; Snippert et al., 2010). While our study is focused on breast cancer modeling, it should be noted that, due to the modular nature of the MADM system and the availability of a genome-wide library of MADM mice (Contreras et al., 2021), one could establish many other cancer models to study their initiation and pre-malignant progression upon the sporadic loss-of-heterozygosity of relevant tumor suppressor genes.

The observed progression trajectory of MADM-mutant mice provides multiple insights for understanding the early genesis of basal-like breast cancer. Although hyper-alveolarization was initially thought to reflect the aberrant expansion of mutant alveolar cells (Tao et al., 2017), our study shows that it actually originates from mutant ductal cells, which likely mis-differentiates toward an alveolar fate upon further progression. In support of this interpretation, recent single-cell sequencing of mammary glands from *Brca1*, *p53*-deficient mice spanning various pre-malignant ages revealed dysregulation of transcription factors driving alveologenesis in luminal progenitor cells, causing aberrant alveolar outgrowth (Bach et al., 2021). Intriguingly, this finding may point to distinct roles of hormonal signaling between pre-malignancy and malignancy of basal-like breast cancer: the hyper-alveolarization of mutant ducts resembles alveolar outgrowth during early pregnancy (Macias and Hinck, 2012), a process known to be regulated by the progesterone signaling (Brisken et al., 1998; Humphreys et al., 1997; Lydon et al., 1995); and progesterone receptors are overexpressed in *Brca1*-deficient mammary epithelial cells of both human and mouse, and exposure to exogenous progesterone dramatically increases mammary gland volume in *Brca1*-deficient mice (King et al., 2004; Ma et al., 2006; Poole et al., 2006). Therefore, although basal-like breast cancers are negative for hormonal receptors at malignancy, the role of progesterone signaling at pre-malignancy warrants further study and may offer a new avenue of cancer prevention for predisposed women (Nolan et al., 2016; Sigl et al., 2016; Trabert et al., 2020).

Upon micro-invasion, MADM-mutant cells undergo a partial luminal-to-basal transition, which could increase the stemness and invasiveness of *Brca1* or *p53*-deficient cells as reported in the literature (Bai et al., 2022; Kim et al., 2011; Liu et al., 2008; Luond et al., 2021; Mizuno et al., 2010) and was recently implicated as a critical step at the onset of basal-like tumorigenesis (Landragin et al., 2022). Our time course analysis of *Brca1*, *p53*-deficient cells in vivo showed that the loss of *Brca1* and *p53* does not immediately induce a luminal-to-basal transition. Instead, mutant cells develop for some time as localized pre-malignancies and only show a partial luminal-to-basal transition at ∼10 months after mutant cells progress to micro-invasive lesions. While we cannot rule out cell-intrinsic mechanisms for this transition, our observation implies that the exposure of luminal cells to extrinsic stromal factors due to basement membrane breaching in micro-invasion, could be the trigger for the luminal-to-basal transition, which we plan to investigate more thoroughly in the future.

While powerful, it takes ∼ 8 months for MADM-mutant mice to progress into the pre-malignant stages described in this study. If desired, additional clinically relevant mutations could be introduced to accelerate cancer development in our model (Annunziato et al., 2019). Compound mutations that are syntenic with *Brca1* and *p53* can be introduced using the same scheme as shown in **Figure S1B**; for mutations that are not syntenic, mutant alleles can be introduced into the MADM model through conventional breeding schemes (Muzumdar et al., 2016; Yao et al., 2020). Another drawback of the current MADM-mutant model is its mixed genetic background that precludes allotransplantation experiments. Therefore, our ongoing effort to backcross these mice into the FVB background should improve the versatility of this model. Notwithstanding these limitations, the current MADM model for basal-like breast cancer enables spatially resolved analysis at any time of pre-malignancy, and can greatly facilitate studies of cancer early detection and prevention.

## Materials and Methods

### Animal

The following mouse lines were crossed to establish the MADM-mutant and MADM-wildtype mice: TG11ML (stock NO. 022976 JAX) (Henner et al., 2013), GT11ML (stock NO. 022977 JAX) (Henner et al., 2013), *Brca1^flox^* (strain NO. 01XC8; NCI) (Xu et al., 1999), *p53^KO^* (stock NO. 002101; JAX) (Jacks et al., 1994), *MMTV-Cre* (stock NO. 003553 JAX) (Wagner et al., 1997). The breeding schemes are shown in Fig. S2. All animal work was performed in the University of Virginia Animal Vivarium. All procedures, including housing and husbandry were approved by the Institutional Animal Care and Use Committee (IACUC) at the University of Virginia, following national guidelines to ensure the humanity of all animal experiments.

### Genotyping

For genotyping, the mouse toe was used to extract DNA for PCR. 120 µl of 50 mM NaOH was added to each toe and then incubated at 95°C for 20 min in the PCR machine, followed by adding 30 µl of 1 M Tris-HCl (pH 7.4) and mixing. 1 µl of toe solution was used for PCR template in a 20 µl PCR reaction. The PCR primer sequences are listed below.

1) *Eif* for MADM TG/GT cassettes: primer-1, 5-TGGAGGAGGACAAACTGGTCAC-3; primer-2, 5-TCAATGGGCGGGGGTCGTT-3; primer-3, 5-TTCCCTTTCTGCTTCATCTTGC-3; PCR products, knock-in (KI) band, 230 bps and WT band, 350 bps. 2) *MMTV-Cre*: primer-1, 5-CACCCTGTTACGTATAGCCG-3; primer-2, 5-GAGTCATCCTTAGCGCCGTA-3; PCR product, KI band, 300 bps. 3) *p53^KO^*: primer-1, 5-ACCGCTATCAGGACATAGCGTT-GG-3; primer-2, 5-CACAGCGTGGTGGTACCTTATG-3; primer-3, 5-GGTATACTCAG-AGCCGGCCTG-3; PCR products, KI band, 700 bps and WT band, 450 bps. 4) *Brca1^flox^*: primer-1, 5-CTGGGTAGTTTGTAAGCATCC-3, primer-2, 5-TCTTATGCCCTCAGAAAACTC-3; PCR products, flox/flox band, 365 bps and WT band, 297 bps.

### Immunofluorescence and immunohistochemistry

Mammary glands were harvested and fixed with 4% paraformaldehyde (PFA) at 4°C for 24 h. For immunofluorescence, tissues were then washed with PBS twice, soaked with 30% sucrose at 4°C for 48 h, and embedded in optimal cutting temperature (OCT). The tissues were sectioned at 20 μm thickness with Thermo NX50 Cryostat. For staining, slides were first blocked in 0.3% Triton-X 100 and 5% normal donkey serum in PBS for 20 min, then incubated with primary antibodies (CK8, Abcam ab182875,1:200. CK14, Biolegend 905301,1:400; E-cadherin, Biolegend 147301,1:200; α-SMA, Sigma A5228, 1:500) diluted in blocking buffer at 4°C overnight. Secondary antibody incubation was performed for 1 h at room temperature in PBS. To stain nuclei, slides were incubated in DAPI solution (1 µg/mL in PBS) for 5 min before being mounted with 70% glycerol. Fluorescent images were acquired on Zeiss LSM 700/710 confocal microscope. Images were processed with Zen and Fiji. For immunohistochemistry, PFA fixed tissue was further processed for paraffin embedding and then sectioned at 4 μm thickness. After antigen retrieval, the primary antibody (Ki67, Epitomics 4203-1, 1:400) was incubated at 4°C overnight, HRP-conjugated secondary antibodies were then used, and 3,3′-diaminobenzidine (Vector Laboratories, SK-4100) was used to develop color.

### Tissue clearing with the CUBIC method and 3D imaging

PFA-fixed mammary glands were cleared for large-scale 3D imaging with the standard CUBIC method (Susaki et al., 2015). Briefly, tissues were immersed in 50% reagent-1 (25 wt% urea, 25 wt% Quadrol, 15 wt% Triton X-100, 35 wt% dH_2_O), shaken at 110rpm 37°C for 12 h, and then transferred to 100% reagent-1 with DAPI (1 µg/ml) for shaking until mammary glands became transparent. After reagent-1, tissues were washed three times with PBS, 1 h each time with shaking to remove the reagent-1. Tissues were then shaken in 50% reagent-2 for 12h at 37°C, followed by 100% reagent-2 (25 wt% urea, 50 wt% sucrose, 10 wt% triethanolamine, 15 wt% dH_2_O), shaking for 48 h. The Zeiss Z.1 light-sheet microscopy system was used for acquiring images. Tissues were attached to the holder of the light-sheet microscope with super glue.

### Gene expression profiling

About 50 mg of tissue from each mammary tumor was used for RNA extraction. The tissue was homogenized in 500 µl TRlzol, then 100 μl chloroform was added and mixed thoroughly, followed by centrifugation (12,000 rcf) for 15 mins at 4°C. The upper layer aqueous phase containing the RNA was transferred to a new tube, and 5 μg of polyacrylamide was added, followed by an equal volume of 70% ethanol. Afterward, 700 μl of the sample was used as the input for RNA isolation using the RNeasy Mini Kit (QIAGEN) according to the manufacturer’s protocol. The quality of RNA samples was evaluated with Bioanalyzer (Agilent Technologies), and samples with an RNA Integrity Number (RIN) > 8 were used for library preparation. Libraries were prepared with a TruSeq Stranded mRNA Library Prep kit (Illumina). Libraries were multiplexed at an equimolar ratio, and 1.3 pM of the multiplexed pool was sequenced on a NextSeq 500 instrument with a NextSeq 500/550 Mid/High Output v2.5 kit (Illumina) to obtain 75-bp paired-end reads. From the sequencing reads, adapters were trimmed using fastq-mcf in the EAutils package (version ea-utils.1.1.2-779) with the following options: -q 10 -t 0.01 -k 0 (quality threshold 10, 0.01% occurrence frequency, no nucleotide skew causing cycle removal). Quality checks were performed using FastQC (version 0.11.8) and MultiQC (version 1.7). Data were aligned to the murine transcriptome (GRCm38.84) using HISATv2 (version 2.1.0) with options for paired-end reads. HISAT read counts were converted to transcripts and normalized to transcripts per million (TPM) using StringTie (version 2.1.6).

### PAM50 extraction and comparison with TCGA datasets

TCGA breast cancer expression data was obtained from the UCSC genome browser (Ciriello et al., 2015). Human orthologs for mouse genes were obtained from the Ensembl biomart in R using the getAttributes function. For both the human and murine datasets, we obtained the intersection of unique orthologs to obtain a set of 14,980 mice to human ortholog genes to evaluate co-expression. To enable cross-species comparisons, we performed a sample-wise column normalization to obtain new transcripts per million (TPM) estimates that accounted for gross differences in gene abundance between species. Hierarchical clustering of PAM50 genes was performed using “pheatmap” in R using Euclidean distance and “ward. D2” linkage.

### Whole exome sequencing and copy number variation analysis

Genomic DNA was prepared from tumors developed in MADM*-*mutant mice and from the tails of the same mice as the control with a DNeasy Blood and Tissue Kit (Qiagen). Whole-exome sequencing at 100x coverage was performed as a contract service with Genewiz. Raw BCL files were converted to fastq files with bcl2fastq v.2.19 and adapter was trimmed with Trimmomatic v.0.38. Trimmed reads were mapped to the mouse reference genome, and somatic variants and copy number variations were called using the Dragen Bio-IT Platform (Illumina) in somatic mode and a panel of normals to remove technical artifacts. The filtered VCF was annotated with Ensembl Variant Effect Predictor (VEP) v95 for the Ensembl transcripts overlapping with the filtered variants. CNVs that passed quality control filters from Dragen were visualized using the GenVisR v.1.16.1 package in R.

### Statistical analysis

Statistical analysis was performed with GraphPad Prism. Bar graphs were presented as the mean[±[standard error of the mean (s.e.m.) unless otherwise annotated in the figure legend. The normality of data distribution was checked with qqplot in R. Depending on whether the data followed a normal distribution, the Student’s t-test or Mann-Whitney *U* test was used as indicated in the figure legends. The Chi-square test or Fisher’s Exact test was used to test frequency distribution as indicated in the figure legends. Statistical significance is noted by “not significant (n.s.)” p[>[0.05, **p[<[0.01, ***p[<[0.001, and ****p[<[0.0001.

## Acknowledgments

We thank Dr. Bing Xu and Xiaoyu Zhao for providing feedback on the manuscript. We also thank Dr. Pat Pramoonjago at the Biorepository and Tissue Research Facility, Sheri Vanhoose at the Research Histology Core, Dr. Stacey Criswell at the Advanced Microscopy Facility, and Shelly Verling at the vivarium for their assistance on the project. We thank Dr. Ammasi Periasamy at the UVA Keck Center for their assistance with the Zeiss Light-sheet Z.1 microscopy system.

## Data availability

The RNA sequencing data of tumors have been deposited in NCBI GEO (https://www.ncbi.nlm.nih.gov/geo/query/acc.cgi?acc=GSE214433; reviewer token: qhmrmcsylxgtlwt). The genomic sequencing data of tumors have been deposited in NCBI SRA (https://dataview.ncbi.nlm.nih.gov/object/PRJNA885219?reviewer=pptilbg30spbh98clp2lh36nbt).

## Author contributions

Conceptualization: J.Z., H.Z.; Methodology: J.Z., S.S., Y.J., E.C., K.A.J., H.Z.; Validation: J.Z., S.S., E.C.; Formal analysis: J.Z., S.S., E.C., K.A.A., K.A.J.; Investigation: J.Z., S.S., Y.J., E.C.; Resources: J.Z., S.S., Y.J., E.C.; Writing - original draft: J.Z., S.S, H.Z.; Writing - review & editing: J.Z., K.A.J., H.Z.; Visualization: J.Z., S.S., K.A.J., H.Z.; Supervision: K.A.J., H.Z.; Project administration: H.Z.; Funding acquisition: K.A.J., H.Z.

---

**Fig. S1. The breeding scheme to build two stock mouse lines for establishing a MADM-based breast cancer mouse model with *Brca1, p53* deficiency**

(A) Location of MADM TG/GT cassettes, *p53*, and *Brca1* on mouse chromosome 11. The physical locations were indicated.

(B) The breeding scheme to incorporate *p53* and *Brca1* mutations into MADM-TG stock and the *MMTV-Cre* transgenes into the MADM-GT stock. Mating between the TG and GT stock produces MADM *p53-Brca1* mice (MADM mutant) and the control MADM wildtype mice.

**Fig. S2. Additional mechanism to generate yellow cells, and MADM labeling specificity in the mammary gland**

(A) Cre-mediated inter-chromosomal recombination could also occur in G1 or post-mitotic cells (G0), which generate dual-colored yellow cells without altering genotype (heterozygous).

(B) MADM-colored cells locate in the CK8+ luminal layer but not in the CK14+ basal layer. Immunofluorescence staining of CK8 and CK14 was performed with sections of mammary glands from MADM-mutant mice at three months old (*n* =3). Scale bar =50 μm.

(C) Quantification of the percentage of MADM-colored cells that are CK8+ (luminal) or CK14+ (basal) from three MADM-mutant mice at three months old (*n* =3). Data are represented as mean ± s.d.

**Fig. S3. The hormone receptor status of MADM tumors and CNVs in human basal-like breast cancer**

(A) Representative images of ER, PR, and Her2 status in three more MADM tumors showed a triple-negative phenotype. Immunohistochemistry staining was performed with tumor sections. Scale bar =100 μm.

(B) Analysis with TCGA datasets for human basal-like breast cancer, showing consistent amplification of *FGFR1, MYC*, loss of *RB1*, and over-expression of *MET*.

**Fig. S4. Hyper-alveologenesis occurred specifically in mutant ducts.**

(A) The pipeline for acquiring large-scale 3D images of mammary ducts with the CUBIC clearing method and light-sheet microscopy.

(B) Whole-mount fluorescence imaging of cleared mammary glands showing distinct morphology of mutant ducts. Mammary glands from 4 MADM-mutant mice at 8 months old mice were assessed. Scale bar =100 μm.

(C) H&E staining of hyper-alveolarized mutant ducts and normal-shape control ducts.

(D) The scheme to count the number of alveoli per 100 μm major ducts to quantify ductal-alveologenesis.

(E) Increased ductal-alveologenesis level of the green mutant ducts compared with the yellow heterozygous ducts (internal control) as shown by the quantification of ductal alveologenesis level from 8-month-old mice (*n*=4). Data are represented as mean ± s.e.m. Mann–Whitney test was used, ***<0.0001.

(F) Examples and criteria for categorizing mutant ducts into different alveologenesis levels.

(G) The distribution of ductal-alveologenesis levels among yellow heterozygous ducts (internal control) and green mutant ducts in glands from 8-month-old mice (*n*=6). 49 yellow control ducts and 23 green mutant ducts were imaged and quantified. A Chi-square test was used. **** p<0.0001.

**Fig. S5. Morphology of mammary ducts in MADM-wildtype mice along aging, and comparison between hyper-alveolarized mutant ducts and age-matched wildtype ducts.**

(A) Upper panel: whole-mount fluorescence imaging of mammary glands from MADM-wildtype mice at a cohort of ages, showing no expansions of GFP+ foci or prominent morphological change of mammary ducts. Scale bar =5 mm. Lower panel: H&E staining showing no morphological changes of mammary ducts along mice aging. Scale bar =200 μm. *n*=3 for each age.

(B) H&E staining of hyper-alveolarized mutant ducts and age-matched wildtype ducts. Pathological review reports that hyper-alveolarized mutant ducts are characterized by nucleomegaly and loss of basal-oriented nuclear polarity. Representative images from four mice. Scale bar =100 μm.

